# Wnt/Beta-Catenin Signaling Is Active in Neuroendocrine Prostate Cancer

**DOI:** 10.1101/2025.03.25.645248

**Authors:** Yingli Shi, Shu Yang, Lin Li, Siyuan Cheng, Jeyaluxmy Sivalingam, Elahe Mahdavian, Xiuping Yu

**Author notes:** Corresponding author’s.

## Abstract

Wnt/beta-Catenin signaling plays a critical role in prostate cancer (PCa) progression, yet its precise contributions in neuroendocrine prostate cancer (NEPCa) remain incompletely understood. In this study, we utilized TRAMP/Wnt-reporter mice to monitor Wnt/beta-Catenin activity and investigated transcriptional alterations associated with NEPCa development. RNA sequencing and pathway enrichment analyses identified neuroactive ligand-receptor interaction, MAPK, calcium, and cAMP signaling as key pathways enriched in NEPCa. Although Wnt signaling was not among the top-enriched pathways, elevated Axin2 expression and increased Wnt-reporter activity suggest its involvement in NEPCa progression. We observed upregulated expression of Wnt3, Wnt6, Dvl2, Dvl3, and Lef1 in NEPCa, coupled with reduced expression of Yap1 and Frat1, which are involved in beta-Catenin degradation. Pharmacological inhibition of Wnt/beta-Catenin signaling using FC101 significantly suppressed PCa growth, underscoring its potential as a therapeutic target. These findings reveal that Wnt/beta-Catenin signaling is active in NEPCa through multiple mechanisms and highlight the need for further investigation into the regulatory interplay between Wnt and YAP1 in prostate cancer.

## INTRODUCTION

Neuroendocrine (NE) differentiation commonly arises in prostate cancer (PCa) following long-term androgen deprivation therapy^1^. Currently, there is no effective treatment for PCa with prominent NE differentiation that presents as “therapy-related” NEPCa^1^. Understanding how NE phenotype emerges subsequent to androgen deprivation therapy is critical for identifying therapeutic targets for advanced PCa.

Accumulating evidence suggests that Wnt/beta-Catenin signaling is activated in advanced PCa^2^. Mutations in this pathway have been detected in 18% of castrate-resistant PCa, and nuclear beta-Catenin, an indicator of active Wnt/beta-Catenin signaling, has been observed in 77% of PCa lymph node and 85% of skeletal metastasis^3^. We have previously shown that Wnt/beta-Catenin signaling is active in advanced human PCa including double-negative PCa (DNPCa) and NEPCa^4,5^. Additionally, we have shown that activation of Wnt/beta-Catenin signaling promotes development of castrate-resistant PCa with an increased NE phenotype, indicating its functional role in castrate-resistant progression and NE differentiation in PCa^6^.

The TRAMP mouse model is widely used for PCa research, where SV40 T-antigen is expressed in the prostate of transgenic mice^7-9^. Similar to human PCa progression, castration accelerates the development of NE tumors in TRAMP mice^7-9^. The Wnt-reporter mouse model used in this study is a transgenic line in which GFP is fused to histone H2B and placed under the control of a Wnt/beta-Catenin responsive promoter^10^. Active Wnt/beta-Catenin signaling induces GFP-histone expression, which is readily detectable in cell nuclei.

In this study, we investigated whether and how Wnt/beta-Catenin is activated in PCa and whether blocking this pathway inhibits NEPCa progression.

## METHODS

### Cell culture

PCa cells were cultured in RPMI 1640 supplemented with 10% heat-inactivated fetal bovine serum and 1% Penicillin-streptomycin. For androgen treatment experiments, cells were cultured in media with 5% charcoal-stripped serum supplemented with ethanol or 10 nM DHT. Cell proliferation was assessed using the IncuCyte method.

### Animal experiments

TRAMP (003135)^11^, Wnt-reporter (013752) ^10^, and Dkk1 (024746)^12^ breeder mice were obtained from Jackson Laboratory and maintained at LSU Health Shreveport animal facility. For the induction of Dkk1 expression, genotype confirmed mice were fed with doxycycline (0.5 mg/ml in water) at 17-18 weeks of age, castrated one week later and sacrificed four weeks post-castration.

### Immunohistochemistry and immunofluorescence staining

Primary antibodies used for immunostaining included GFP (Cell Signaling), FOXA2 (Abcam)), and T-antigen (Santa Cruz). IHC staining was performed using the Vectastain elite ABC peroxidase kit (Vector Laboratories, Burlingame, CA).

### RNA sequencing

Paired-end RNA sequencing was conducted by Novogene. Differentially expressed (DE) genes were identified using DESeq2, with a significance threshold of padj < 0.05 and an absolute log2 fold change (|log2FC|) > 1. Functional enrichment analysis was performed using Gene Ontology (GO) and Kyoto Encyclopedia of Genes and Genomes (KEGG) pathway analyses. Enrichment analysis and visualization of enrichment results were conducted using the “clusterProfiler” package in RStudio.

## RESULTS

### Wnt/beta-Catenin signaling is active in TRAMP NEPCa

To evaluate Wnt/beta-Catenin signaling activity in NEPCa, we established TRAMP/Wnt-reporter bi-transgenic mice. Immunohistochemistry staining for GFP revealed that 21 out of 24 NEPCa tumors expressed GFP, concurrent with T-antigen and the NEPCa marker FOXA2, indicating active Wnt/beta-Catenin signaling (Figs. 1A-F). However, a subset of NEPCa tumors (n=3) was GFP-negative (Figs. 1G-I).

**Fig. 1:**
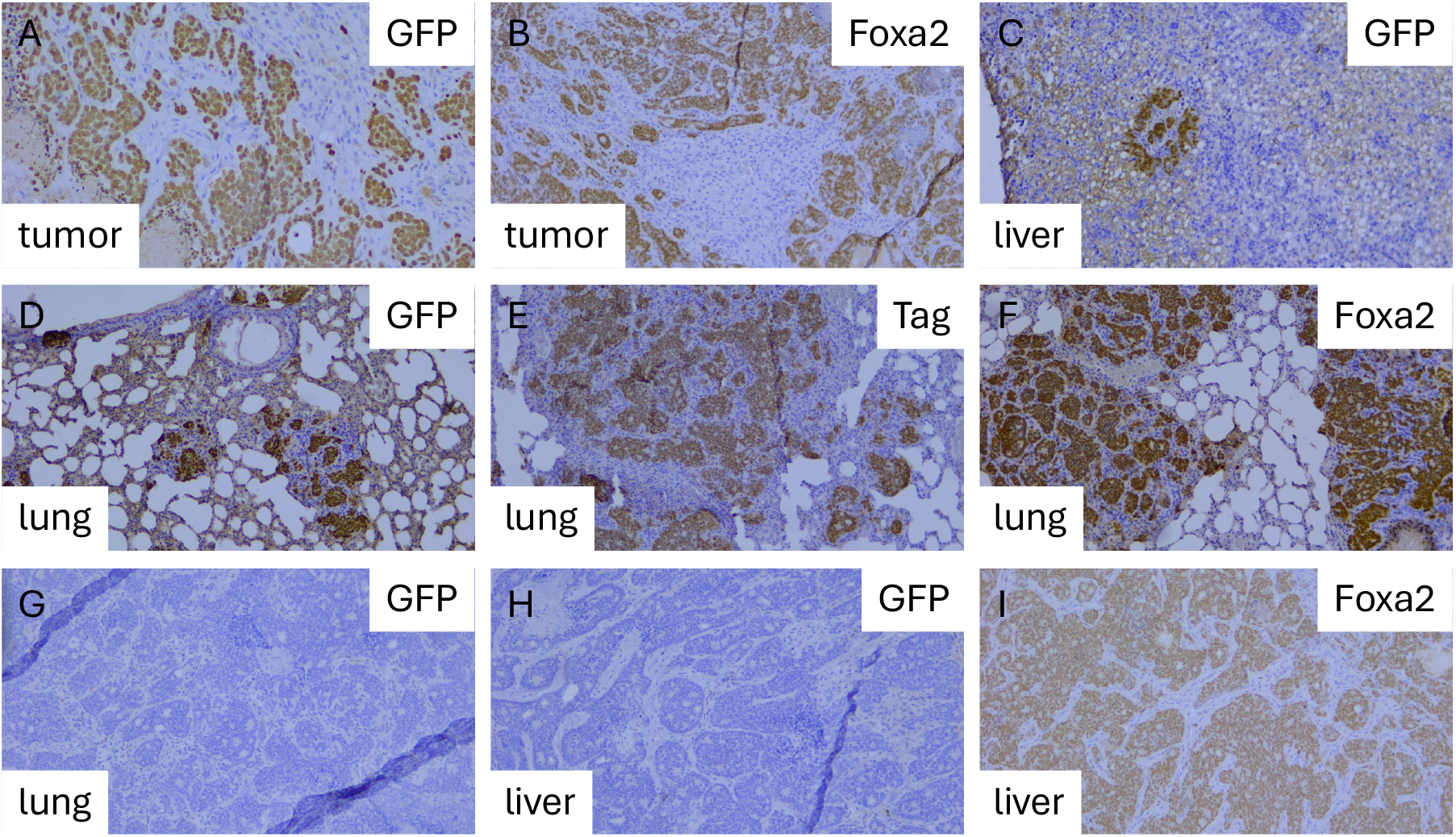
Immunohistochemical staining of TRAMP/Wnt-reporter tumors. (A-F) Representative images of IHC staining showing GFP expression in tumors, indicating active Wnt/beta-Catenin signaling. (G-I) IHC staining of tumors lacking GFP expression.

RNA sequencing and pathway enrichment analysis of TRAMP tumors with or without NEPCa (Figs. 2A-B) identified neuroactive ligand-receptor interaction, MAPK, calcium, and cAMP signaling as the top enriched pathways in NEPCa. While Wnt signaling was not among the enriched pathways, the expression of Axin2, a direct downstream target of Wnt/beta-Catenin signaling, was elevated in NEPCa (Fig. 2C), aligning with the increased Wnt activity observed in TRAMP/Wnt-reporter mice. Further analysis revealed increased expression of Wnt3, Wnt6, Lrig1, Fzd3, Dvl2, Dvl3 and Lef1 in NEPCa, while Yap1 and Frat1, key regulators of beta-Catenin degradation, was downregulated (Fig. 2C). In contrast, multiple Wnt receptors and co-receptors were highly expressed in AdPCa tumors (Fig. 2C). Given the higher Wnt-reporter expression observed in NEPCa compared to AdPCa cancer cells, it is possible that these receptors and co-receptors are predominantly expressed in other cell types within the tumor microenvironment.

**Fig. 2:**
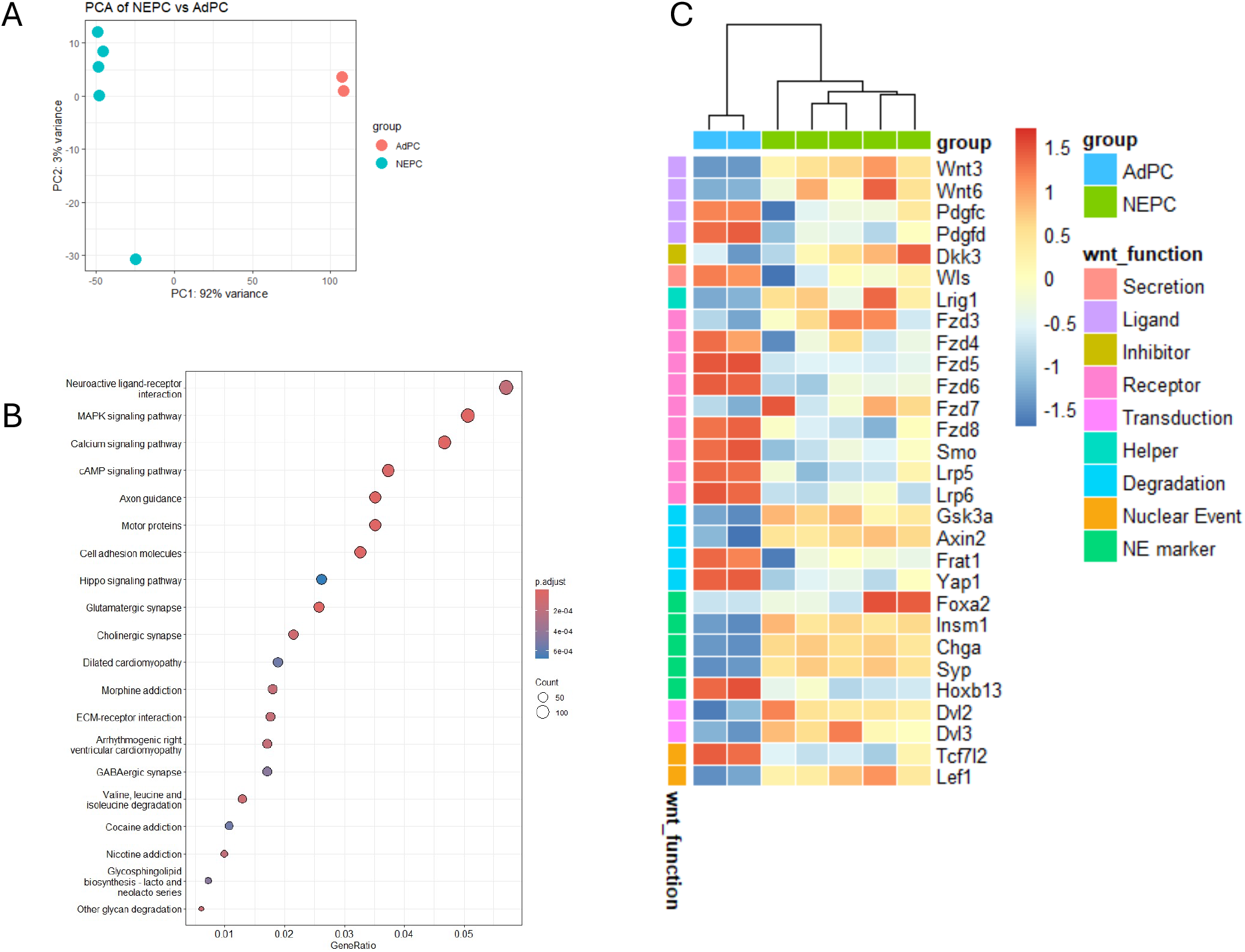
RNA sequencing analysis of TRAMP NEPCa versus AdPCa. (A) Principal component analysis (PCA) illustrating transcriptomic differences between NEPCa and AdPCa. (B) Gene Ontology (GO) enrichment analysis highlighting pathways significantly altered in NEPCa. (C) Heatmap displaying differentially expressed NEPCa markers and Wnt-related genes in NEPCa compared to AdPCa.

### Wnt/beta-Catenin signaling is active in advanced human PCa

Analysis of Wnt/beta-Catenin signaling in human PCa using the HuPSA single-cell RNA sequencing dataset indicates that Wnt/beta-Catenin activity is most prominent in dendritic cells^4^. In PCa cells, Wnt activity is high in double-negative PCa (DNPCa) and NEPCa^5^. However, examination of Wnt-related gene expression showed minimal expression of canonical Wnt ligands and receptors in PCa cells (Fig. 3). But consistent with TRAMP NEPCa findings, YAP1 and WWTR1 (encoding TAZ) expression was low in human NEPCa cells (Fig. 3).

**Fig. 3:**
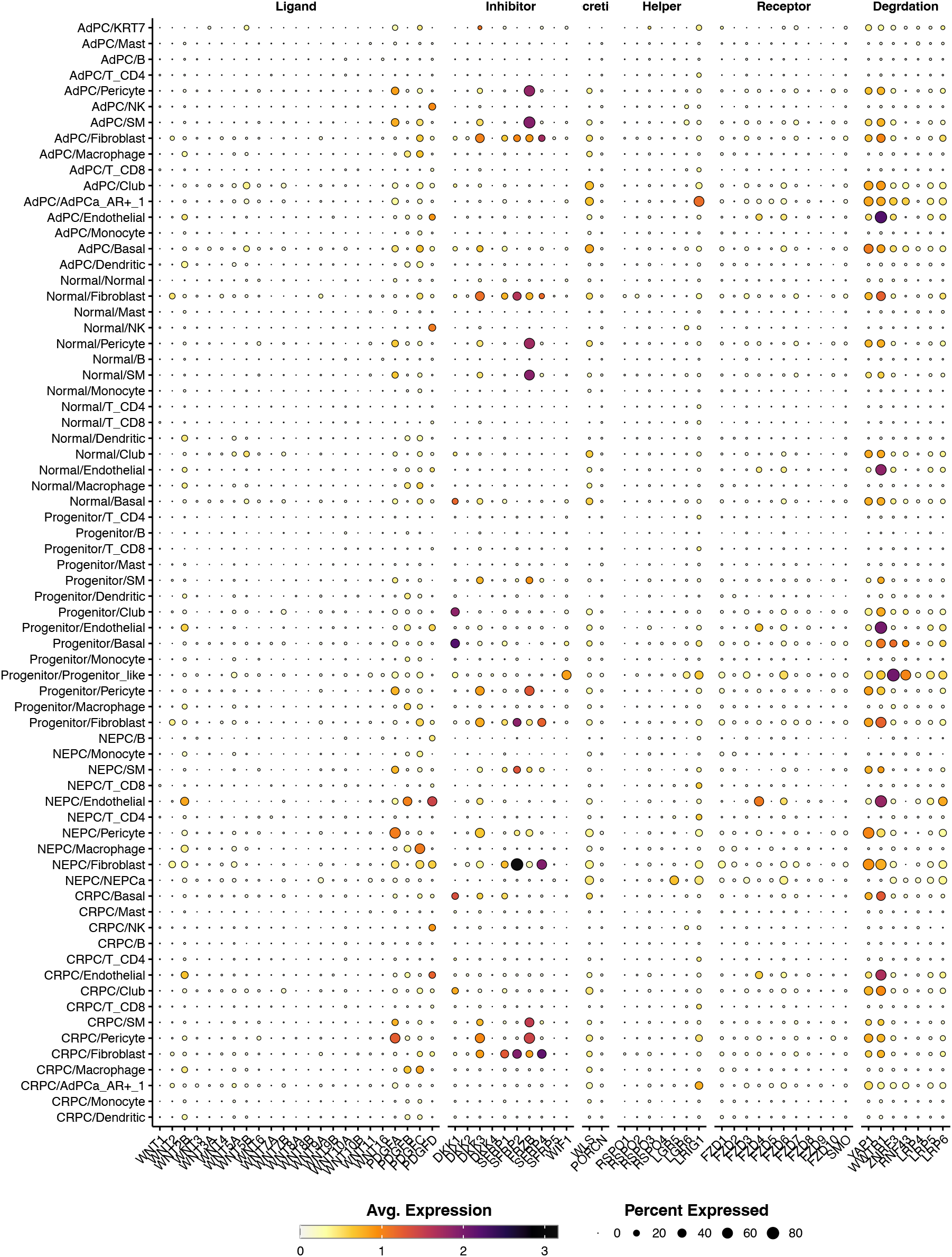
Expression of Wnt pathway components across various cell types in human PCa. PCa cells exhibit minimal expression of canonical Wnt ligands and receptors. Notably, YAP1 is highly expressed in prostate adenocarcinoma but significantly downregulated in NEPCa. A similar pattern is observed for the WWTR1 gene, encoding TAZ, another key transcription factor in the Hippo signaling pathway.

### Overexpression of DKK1 reduced NEPCa incidence

Dkk1 is a secretory inhibitory factor of canonical Wnt signaling. To determine whether blocking Wnt/beta-Catenin signaling prevents NEPCa progression, we ectopically expressed Dkk1 in TRAMP tumors using inducible Dkk1 transgenic mice^12^. We found that overexpression of Dkk1 reduced NEPCa incidence in TRAMP tumors (29% in TRAMP vs 16.7% in TRAMP/Dkk1, Chi-square test, p = 0.1849); however, but the reduction was not statistically significant.

### Blocking Wnt/beta-Catenin signaling inhibits PCa growth

To assess the role of Wnt/beta-Catenin activation in therapy resistance, PCa 22Rv1 cells were treated with 5 nM paclitaxel in the presence or absence of Wnt3A (50 ng/mL). We found that Wnt3A did not significantly alter cell proliferation (Fig. 4A).

**Fig. 4:**
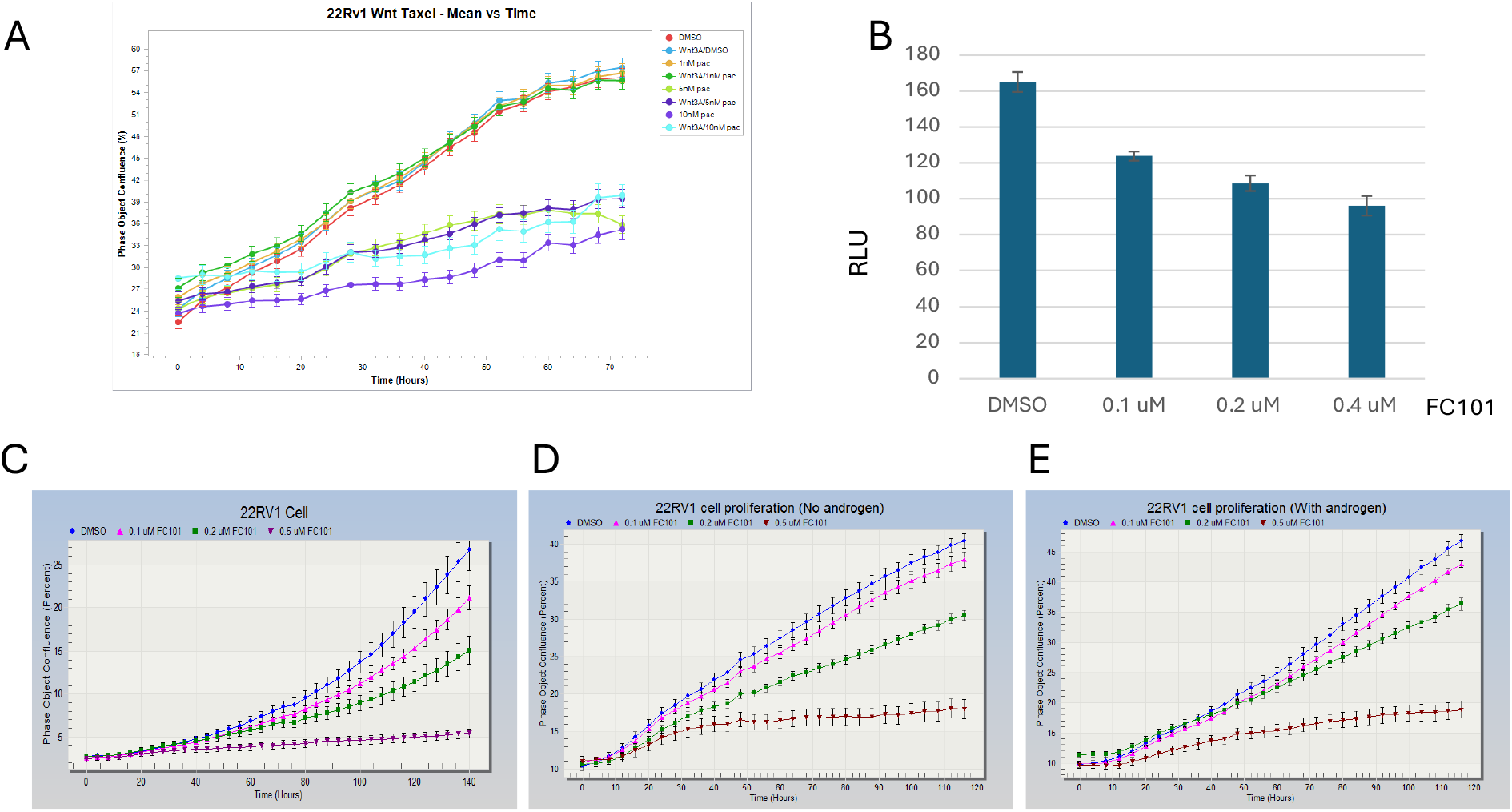
Cell proliferation assays evaluating Wnt/beta-Catenin signaling and therapeutic response. (A) IncuCyte proliferation assay of 22Rv1 PCa cells treated with paclitaxel (1, 5, or 10 nM) in the presence or absence of WNT3A. The addition of WNT3A did not significantly affect the proliferation rates of paclitaxel-treated cells. (B) Luciferase reporter assay in PC3 cells transfected with the TOP-flash Wnt-reporter plasmid, demonstrating dose-dependent inhibition of Wnt/beta-Catenin signaling by FC101. (C-E) IncuCyte proliferation assays evaluating FC101’s effect on PCa cell growth under different culture conditions: (C) complete media, (D) charcoal-stripped media (androgen-deprived), and (E) charcoal-stripped media supplemented with androgens. FC101 inhibited PCa cell proliferation under both androgen-sufficient and androgen-deprived conditions.

To test whether pharmacologically blocking Wnt/beta-Catenin signaling suppresses PCa growth, we blocked Wnt signaling using FC101^14^, a molecule that inhibit Wnt/beta-Catenin signaling in PCa cells (Fig. 4B). We found that FC101 suppressed PCa cell growth under both androgen-sufficient and deficient conditions (Figs. 4C-E).

## DISCUSSION

In this study, we used TRAMP/Wnt-reporter mice to directly monitor the activity of Wnt/beta-Catenin signaling in PCa. Our findings indicate that Wnt/beta-Catenin signaling is activated in advanced PCa through multiple mechanisms, including contributions from the tumor microenvironment, increased expression of canonical Wnt ligands and transcription factors, and reduced expression of the factors involved in beta-catenin degradation. Notably, although Wnt ligands are expressed at low levels, their effects can be amplified by the loss of YAP1^15^. Given the established role of YAP1 in regulating cellular differentiation and maintaining epithelial identity, its downregulation in NEPCa suggests a potential shift toward a more plastic, undifferentiated state. The precise mechanisms governing YAP1 suppression in NEPCa remain unclear and warrant further investigation.

The interpretation of bulk RNA sequencing data presents certain limitations, as it reflects an average measurement of Wnt activity across prostatic tumors. This averaging effect may obscure enrichment signals due to the presence of stromal and immune cell populations within the tumor microenvironment. However, the observed elevated expression of Axin2, a well-established Wnt target gene, supports the conclusion that Wnt/beta-Catenin signaling is indeed active in NEPCa. This finding is consistent with previous reports linking Wnt activation to aggressive, castration-resistant prostate cancer subtypes.

Interestingly, the presence of GFP-negative NEPCa tumors in our reporter model suggests that alternative mechanisms may be driving NE differentiation independent of Wnt/beta-Catenin activation. This observation raises the possibility of heterogeneous signaling pathways contributing to lineage plasticity in advanced PCa. Future studies employing single-cell transcriptomics and lineage-tracing approaches could provide deeper insights into the cellular heterogeneity and regulatory networks underlying NEPCa progression.

Additionally, our findings have therapeutic implications. Given the complexity of Wnt/beta-Catenin regulation in PCa, targeting this pathway in advanced disease requires a nuanced approach. Strategies that selectively modulate pathway components while considering the tumor microenvironment and YAP1-associated signaling could offer novel therapeutic opportunities. Furthermore, the interplay between Wnt signaling and immune regulation remains an area of interest, as recent evidence suggests that Wnt activation may influence tumor immune evasion mechanisms^16^. Investigating these interactions could provide valuable insights into potential combinatorial treatment strategies.

## ACKNOWLEDGEMENT

This research was supported by NIH R01 CA226285 and LSU Health Shreveport FWCC Stimulus grants to X. Yu, LSU Health Shreveport FWCC Carroll Feist postdoctoral Fellowship to S. Cheng, and LSU Health Shreveport FWCC Carroll Feist predoctoral Fellowship to L. Li.

